# Female Alms1-deficient mice develop echocardiographic features of adult but not infantile Alström Syndrome cardiomyopathy

**DOI:** 10.1101/2023.10.16.562570

**Authors:** Eleanor J. McKay, Ineke Luijten, Adrian Thomson, Xiong Weng, Katya Gehmlich, Gillian A. Gray, Robert K. Semple

## Abstract

**Background:** Alström Syndrome (AS), a multisystem disorder caused by biallelic *ALMS1* mutations, features major cardiac complications often causing early mortality. These are biphasic, including infantile dilated cardiomyopathy, and distinct adult-onset cardiomyopathy. Cardiomyocyte maturation defects, cardiac fibrosis and early atherosclerosis have all been invoked as contributors to heart failure in AS, but their relative importance and inter-relationships are unknown.

**Methods:** Cardiac function of global *Alms1* knockout mice was assessed by echocardiography at postnatal day 15 (P15) and at 8 and 23 weeks of age. Echocardiography was also undertaken in female mice with *Pdgfrα*-Cre-driven *Alms1* deletion in cardiac fibroblasts and a small proportion of cardiomyocytes. Histological and transcriptional analysis of myocardium at P15 and 24 weeks of age was also performed.

**Results:** Cardiac function was unaltered in knockout mice of both sexes at P15 and 8 weeks of age. At 23 weeks of age female but not male knockout mice showed increased left atrial area, decreased isovolumic relaxation time, and reduced ejection fraction, consistent with early restrictive cardiomyopathy. No histological or transcriptional changes could be identified in myocardium of 23-week old female *Alms1* KO mice, however. *Pdgfrα*-Cre-driven *Alms1* KO in females did not recapitulate the phenotype of global KO at 23 weeks.

**Conclusions:** Adult female, but not male, *Alms1*-deficient mice show echocardiographic evidence of cardiac dysfunction, consistent with the restrictive cardiomyopathy of AS. The explanation for sexual dimorphism remains unclear, but may involve metabolic or endocrine differences between sexes. No infantile cardiomyopathy was found in this study.

## Introduction

Alström syndrome (AS) is an autosomal recessive disorder caused by biallelic loss-of-function mutations in the *ALMS1* gene. The product of the *ALMS1* gene is a large, 460 kDa protein primarily localised to the centrosome and basal body of primary cilia. In keeping with this, cardinal features of AS include infantile rod-cone retinal dystrophy, sensorineural deafness, obesity, and diabetes mellitus, which are common features of several so called primary “ciliopathies”. AS also features prominent cardiac complications, which are a major cause of early morbidity and mortality in the syndrome^1^.

Cardiac dysfunction occurs in approximately 60% of patients with AS at some point. The natural history of cardiac manifestations of AS is complicated, however, with a biphasic pattern across the lifecourse. Dilated cardiomyopathy and congestive heart failure occurs in 43% of infants with AS, usually reported in the first 12 months of life. In some cases this is fatal, but with treatment around three quarters of patients recover within 3 years^2^. The mechanism of infantile cardiac dysfunction in AS is poorly understood. One clue was offered by identification of biallelic *ALMS1* mutations in four infants who died or underwent heart transplantation due to mitogenic cardiomyopathy, defined by persistent postnatal mitogenesis of cardiomyocytes^3,4^. Whether this was a sentinel presentation of AS is unknown, but similar histological appearances were reported in *Alms1*^GT/GT^ mice harbouring a gene trap in intron 13, resulting in global knockout (KO) of *Alms1*^3^. This remains the clearest potential explanation for the infantile heart failure of AS to date, but corroboratory studies are needed. How loss of ALMS1 expression might induce persistent mitogenesis is unknown.

Around 30% of adults with AS develop heart failure, in half of whom this is fatal or leads to transplantation^2,5^. This appears independent of infantile cardiomyopathy (13% of patients develop both infantile and adult-onset cardiac dysfunction and 18% of patients only develop adult-onset dysfunction)^2,6,7^. Adult-onset cardiomyopathy in AS features myocardial hypertrophy and dilation with progressive fibrosis and restrictive impairment of both ventricles^1^. Accelerated atherosclerosis and coronary artery disease (CAD) are also reported^7–10^. Duration of diabetes in AS is predictive for aortic pulse wave velocity, and thus cardiovascular events^10^, but no association between CAD and cardiac fibrosis was found ^7^. 63% of patients with AS develop chronic kidney disease (CKD)^11^, and 30% have hypertension^2,12^, both further possible drivers of CAD.

Cardiac pathology in AS may thus be a composite result of cardiomyocyte-autonomous developmental defect in infancy, with accelerated atherosclerosis, and progressive fibrosis becoming prominent in young adulthood, driven in part by exogenous factors such as diabetes, insulin resistance, dyslipidaemia, hypertension and impaired renal function. Unpicking the relative contributions of these processes to the strikingly poor cardiovascular outcomes in AS is difficult or impossible in human studies where they are generally admixed. This issue is highly important as it will guide choice of experimental treatments for future trials in AS. For example, the high prevalence of cardiac fibrosis suggests potential value for anti-fibrotic therapies^13^, however whether fibrosis is a cause of cardiac dysfunction or an epiphenomenon arising from a chronic repair process in AS is unknown.

Understanding the cardiac phenotype of *Alms1* KO mice may be clinically relevant not only to patients with AS, but may also provide insights into common cardiac disease. In adults with AS, heart failure occurs alongside obesity, severe insulin resistance and shows preserved ejection fraction^14^, similar to heart failure with preserved ejection fraction (HFpEF) for which metabolic syndrome is a major risk factor.

Several *Alms1* KO mouse lines have been described. These recapitulate many key features of AS, including vision and hearing loss, obesity and insulin resistant diabetes^15–17^. Surprisingly, given its importance to patients with AS, no detailed evaluation of cardiac function in murine AS models has been offered to date. We thus set out to interrogate cardiac phenotypes across the lifecourse in a new *Alms1* global KO mouse model. Furthermore, to help distinguish heart autonomous and non-autonomous drivers of cardiac complications, we also studied mice with *Alms1* KO only in mesenchymal stem cells and their descendants using the constitutive *Pdgfrα*-Cre driver. This has been shown to drive recombination in cardiac fibroadipogenic precursor cells, such as cardiac fibroblasts, but only in 18% of cardiomyocytes^18^. *Pdgfrα*-Cre also drives recombination in preadipocytes and their descendants, in several brain regions^19^, but not in liver or muscle^20–22^. All mice were fed *ad libitum* with obesogenic high fat diet from six weeks of age to exacerbate systemic IR and any propensity to HFpEF, and to allow correlation with prior detailed metabolic studies undertaken on the same diet^19^.

## Materials and Methods

### Mouse strains and experimental protocols

*Alms1*^tm1c(EUCOMM)Hmgu^ mice with loxP sites flanking exon 7 were generated by and purchased from GenOway, France. Excision of the floxed *Alms1* exon 7 yields a premature stop codon in exon 8. CAG-Cre mice^23^ were a gift from Dr Matthew Brook and were used to generate a global *Alms1* KO. *Pdgfrα*-Cre mice^20^ were purchased from The Jackson Laboratory (Strain #013148) and used to KO *Alms1* in mesenchymal stem cells. Although no reliable antibody against murine Alms1 was available to prove absence of Alms1 protein in the KO mice generated, recombination and loss of exon 7 of the *Alms1* transcript was confirmed both in genomic DNA and in cDNA from heart (**Figure S1A**), liver and adipose tissue by qPCR^19^, as described in Supplementary Material.

All mice were purchased and maintained on a C57/BL6/N background, confirmed by single nucleotide polymorphism profiling (Transnetyx). Genotyping was performed by Transnetyx using real-time PCR in all cases except neonatal studies in which genotyping was performed in-house. In-house genotyping was performed by DNA gel electrophoresis following PCR amplification. One common forward primer (ATACCACCACACCTGGGAGG) and two reverse primers were used: reverse 1 (sequence contained in loxP flanked region; CACCATGTAAACACTAGAAATAGAACCCAGGTC) and reverse 2 (GCCAGGAGGAGCAAGACAAT). Presence of WT *Alms1* results in generation of a 366bp fragment by forward and reverse 1 primers. Excision of the loxP flanked sites results in the generation of a 306bp fragment by forward and reverse 2 primers.

Mice were group-housed in individually ventilated cages (IVCs) at the biological research facility at the University of Edinburgh. A 12-hour light/dark cycle (lights on at 0700 and off at 1900) and controlled temperature/humidity (19-21°C/50%) were maintained. Until 6 weeks old, mice had *ad libitum* access to standard chow (CRM, Special Diet Service), then replaced by 45% fat diet (D12451, Research Diets). Mice were single housed from 13 weeks of age. All experimental protocols were approved by the University of Edinburgh Biological Science Services and performed in compliance with the UK Home Office Scientific Procedure (Animals) Act 1983. Echocardiography was performed by a single experienced operator at Edinburgh Preclinical Imaging. Histological and transcriptional analysis is described in Supplementary Material.

### Echocardiographic studies

For echocardiography mice were anaesthetised with 4% isoflurane in 100% oxygen for induction, with maintenance using 1-2% isoflurane. Animals were placed supine on a heated imaging table with paws attached to electrocardiogram (ECG) electrodes. Heart rate was maintained at 450-550 beats per minute (bpm). Body temperature measured via rectal probe was maintained at 37±0.5°C using heated table and a heating lamp. Chest hair was removed using depilatory cream (Nair hair removal cream, Church & Dwight) and ultrasonography was performed using a VisualSonics Vevo 3100 high frequency ultrasound imaging system (FUJIFILM VisualSonics). An MX550D transducer was used except for obese mice at 23 weeks, where a lower frequency MX250 transducer with increased scan depth was required for Doppler measurements.

For left ventricle (LV) function assessment, electrocardiogram-gated Kilohertz visualisation (EKV^24^) was applied on the parasternal long-axis (PSLA) view and for M-mode on the parasternal short axis (PSAX) view at LV midpoint with papillary muscles at 2 and 4 o’clock. Left atrium measurements were performed on EKV-modified right PSLA views. Doppler measurements of LV inflow and outflow were obtained in the apical 4 chamber view. Analysis of echocardiographic data was undertaken using Visualsonics Vevo LAB 5.71 software (FUJIFILM VisualSonics) as detailed in Supplementary Material.

For normalisation of size-dependent measurements (left ventricle mass, ventricle and atria areas), the nose-anus length and tibia length were measured using a digital calliper. Nose-anus length was measured when animals were anaesthetised for echocardiography at 23 weeks of age. Tibia length was measured following dissection at 24 weeks of age. As tibia length proved shorter in male and female global *Alms1* KO animals (**Figure S3A**), nose-anus length, which did not differ by genotype (**Figure S3B**), was used for normalisation. For normalisation of neonatal echocardiography, body mass was used, which did not differ between experimental groups.

### Blinding and statistical analysis

Experimenter and analyser were blinded to genotype wherever possible, including for all *in vivo* studies, tissue processing, RNA extraction, imaging and analysis, and unblinding was automated by an Excel template to prevent accidental memorisation of genotypes. Statistical analysis was performed in GraphPad Prism 9.2.0. A normal distribution was assumed for all data, as the small numbers required by best practice in animal research preclude reliable testing for normality. Student’s t-test was used for comparison of 2 groups and ANOVA for comparison of more than 2 groups. The Bonferroni correction was applied when multiple t-tests were performed. Šídák’s multiple comparisons test was applied following ANOVA to data generated from the same animals at more than one time point, and Tukey’s multiple comparison test following ANOVA to compare values between multiple groups. Linear regression was used to compare two experimental groups where the variable of interest depended on another variable that differed between groups (e.g. heart mass and body mass).

## Results

We first set out to assess whether our novel global *Alms1* KO mouse model recapitulated the infantile cardiomyopathy of AS. One global *Alms1* KO mouse harbouring a gene trap in *Alms1* intron 13 has previously been reported to show persistent mitogenesis in the heart and increased heart to body mass ratio at postnatal day 15 (p15)^3^, a timepoint roughly equivalent to the developmental age at human birth. This is in keeping with the mitogenic cardiomyopathy described in 4 infants with biallelic *ALMS1* loss-of-function mutations^3^. We thus set out to seek corroboratory evidence for this in our newly generated global *Alms1* KO mouse, while extending the prior studies by characterising cardiac function by echocardiography. We elected to undertake functional studies at postnatal day 15 (p15) to replicate the prior study, and because this corresponds to an age of high prevalence of infantile cardiomyopathy in AS. In contrast to the previous study, no changes in heart to body mass ratio were observed in either male or female *Alms1* KO mice at p15 (**Figure 1A,B**). Moreover the number of cardiomyocytes staining for the proliferative marker phosphorylated histone H3 (pH3), previously reported to be increased in myocardium of *Alms1* KO mice at p15^3^, did not differ between *Alms1* KO and WT littermates of either sex (**Figure 1C,D**). Finally, no changes in a range of cardiac anatomical and functional measures obtained echocardiographically were seen. These included left ventricular mass, wall thickness and chamber dimensions (**Figure 1E-G, Figure S1B,C**), measures of systolic and diastolic function, longitudinal strain and ventricular dyssynchrony (Bilchick et al., 2006) (**Figure 1H-J, Figure S1D-H**).

**Figure 1.**
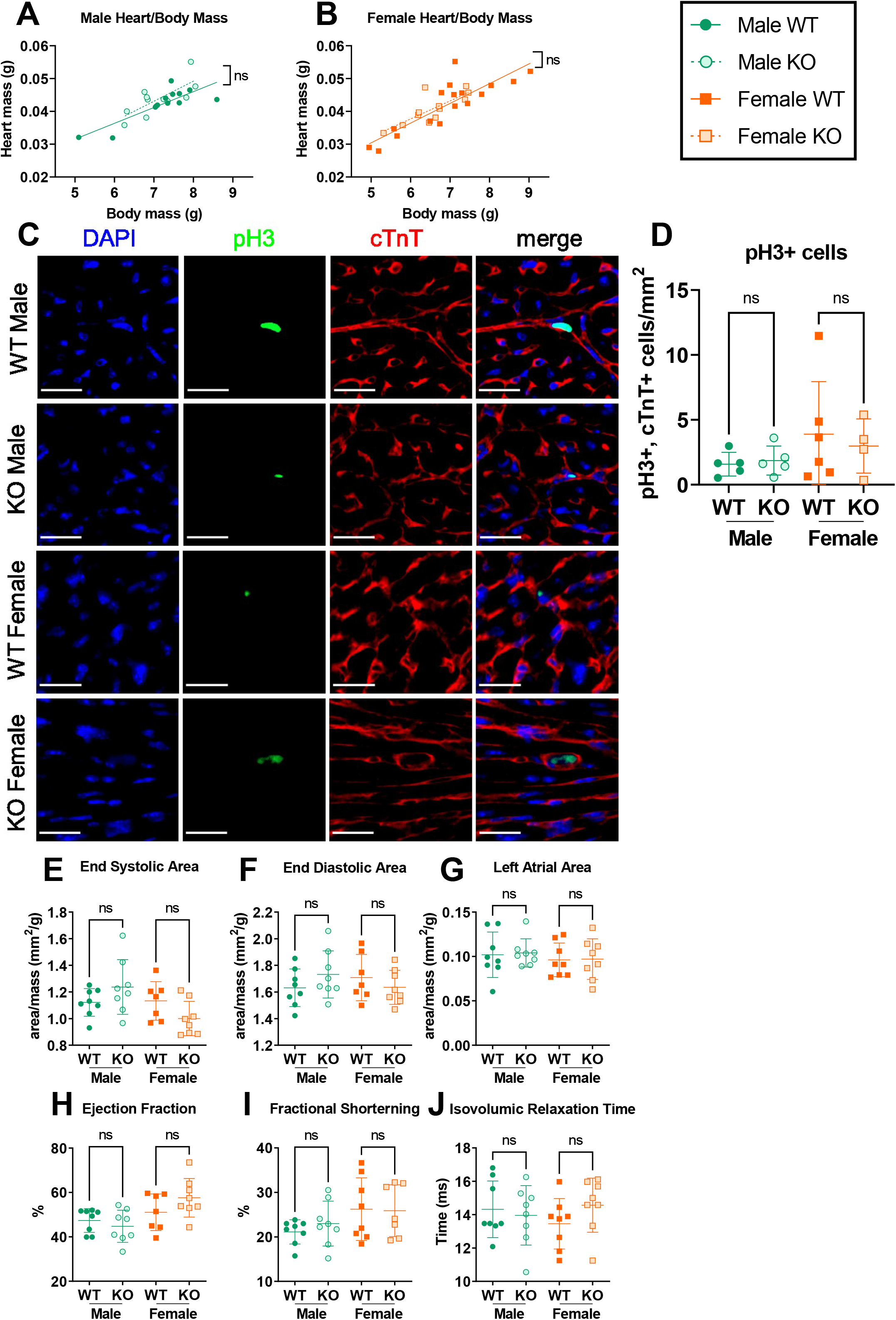
Neither male nor female global *Alms1* knockout mice exhibit a cardiac phenotype at post-natal day 15. (**A,B**) Linear regression of heart and body mass for males and females respectively. (**C**) Representative images of immunofluorescent staining of cardiac left ventricle for proliferative marker phosphorylated histone H3 (pH3) with co-staining for cardiac troponin (cTnT) and DAPI. Scale bars 20μm. (**D**) Quantification of (**C**). (**E-J**) Echocardiography data. Area values calculated from echocardiography (**E-G**) are normalised to total body mass. Each data point represents an individual animal with bars in (**D-J**) representing mean ± sd. Comparison between groups in (**D-J**) was undertaken using two-way ANOVA with Tukey’s multiple comparisons test. Lines in linear regression graphs (**A,B**) represent lines of best fit. Comparisons between lines of best fit was undertaken by simple linear regression, with square brackets showing comparison of y intercepts. No significant change was seen between gradients. For (**A,B**) N = 13, 11, 17 and 12 for WT males, KO males, WT females and KO females respectively. For (**D**) N = 5, 5, 6 and 4 for WT males, KO males, WT females and KO females respectively. For (**E-J**) N = 8, 8, 7 and 8 for WT males, KO males, WT females and KO females respectively.

Consistent with the normal echocardiographic and histological appearances of hearts at P15, transcriptional analysis of myocardium for a panel of markers of heart failure, namely actin alpha 1 (*Acta1)*; myosin heavy chain beta (*Myh7)*; and natriuretic peptides A and B, (*Nppa* and *Nppb)* (**Figure S2A-D**), normalised to *Gapdh* (e.g. **Figure S2E**), showed no clear indications of cardiac dysfunction. An isolated increase in *Myh7* expression between female WT and *Alms1* KO mice is not consistent with expression of the other genes and is of uncertain importance. Collectively these findings provide no evidence of mitogenic or other forms of infantile cardiomyopathy in the new global *Alms1* KO mouse strain.

We next sought to assess cardiac function in global *Alms1* KO mice in adulthood, at both 8 weeks and 23 weeks of age, with tissue collection at 24 weeks of age. These ages correspond roughly to the second and fourth decades of human life, periods in which AS cardiomyopathy is commonly seen. Both male (**Figure 2A**) and female (**Figure 2C**) absolute heart masses were increased at 24 weeks of age, however linear regression showed this increase to be proportionate to body mass, which is higher in AS and *Alms1* KO mice. No changes in echocardiographic anatomical or functional indices were observed in either male or female mice at 8 weeks of age (**Figure 2B,D-L, Figure S3C-R**). At 23 weeks of age, however, the left atrial area of female mice was increased compared to WT littermate controls (**Figure 2D**), but this was not observed in male mice. Increased left atria area is a well established indirect indicator of diastolic cardiac dysfunction. Female global *Alms1* KO mice at 23 weeks also showed several other cardiac functional changes suggestive of systolic dysfunction compared to WT littermate controls, including reduced ejection fraction, fractional ventricular area change and isovolumic relaxation time (**Figure 2G,H,L**). Fractional shortening showed a trend towards reduction (**Figure 2K**). Again, all these indices were unchanged in male mice (**Figure 2B,E,F,I,J**). Left ventricle mass, wall thickness, cross-sectional areas, performance index, longitudinal strain and dyssynchrony were unchanged in both sexes at 23 weeks of age (**Figure S3C-R**).

**Figure 2.**
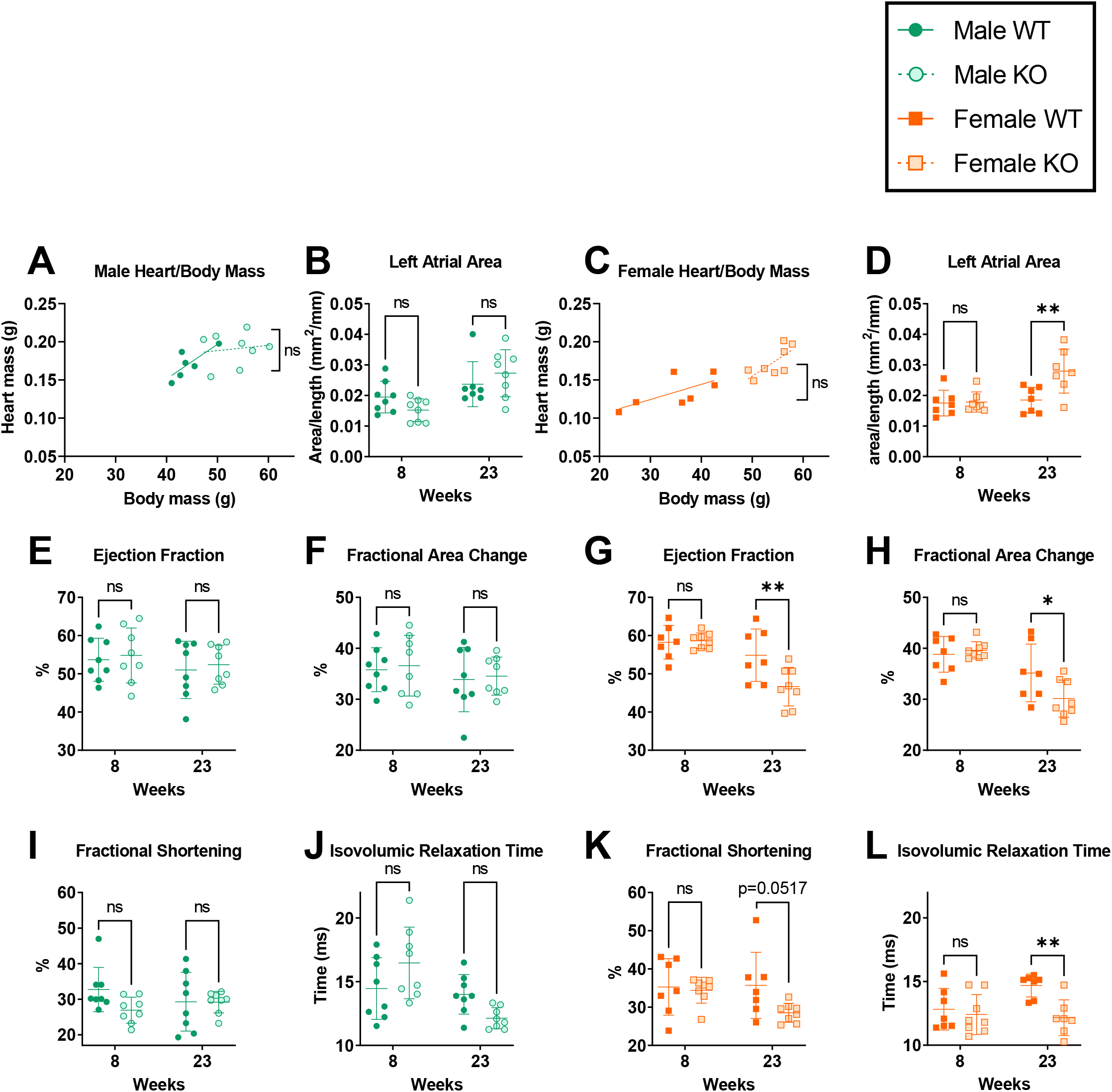
Systolic and diastolic dysfunction develops in female but not male global *Alms1* knockout mice with age. Linear regression of heart to body mass of males (**A**) and females (**C**) at 24 weeks of age. (**B, D-L**) Echocardiography parameters of male and female mice at 8 and 23 weeks of age. Each data point represents an individual animal. Left atrial area values (**B,D**) are normalised to nose-anus length. Bars in (**B, D-L**) representing mean ± standard deviation. Lines in linear regression graphs (**A,C**) represent lines of best fit. Comparison between groups (**B, D-L**) was undertaken using a two-way ANOVA with Šídák’s multiple comparisons test. Comparisons between lines of best fit (**A,C**) were undertaken by simple linear regression, with square brackets showing comparison of y intercepts. No significant change was seen between gradients. * denotes p<0.05, ** denotes p<0.01 and **** denotes p<0.0001. N = 8, 8, 7 and 8 for WT males, KO males, WT females and KO females respectively

Transcriptional analysis of hearts at 24 weeks of age was next undertaken to evaluate markers of cardiomyopathy, as before, with addition of two genes associated with fibrosis, alpha-1 type I collagen (*Col1a1)* and lysyl oxidase (*Lox)*, and three genes involved in cell cycle regulation and cellular senescence, cyclin dependent kinase inhibitor 1A (*Cdkn1a*), cyclin dependent kinase inhibitor 2A (*Cdkn2a*) and lamin B1 (*Lmnb1)*. No changes in expression of any of these genes were seen between female global *Alms1* KO mice and WT littermates (**Figure 3A-I**) when normalised to Gapdh (e.g. **Figure 3J**). Global *Alms1* KO male mice did show decreased transcript levels of *Lmnb1* (**Figure 3I**). However although reduced *Lmnb1* expression is one marker of cellular senescence, *Cdkn1a* and *Cdkn2a* transcripts, which are typically elevated in senescence, were decreased or showed a trend in this direction (p=0.1536 for *Cdkn1a*), arguing against established senescence (**Figure 3G,H**). There were no changes in genes typically associated with heart failure or fibrosis in the male *Alms1* KO mice (**Figure 3A-F**). In keeping with unchanged transcript levels for fibrosis-associated genes, no increase in picrosirius red staining for fibrosis was seen (**Figure 3K**).

**Figure 3.**
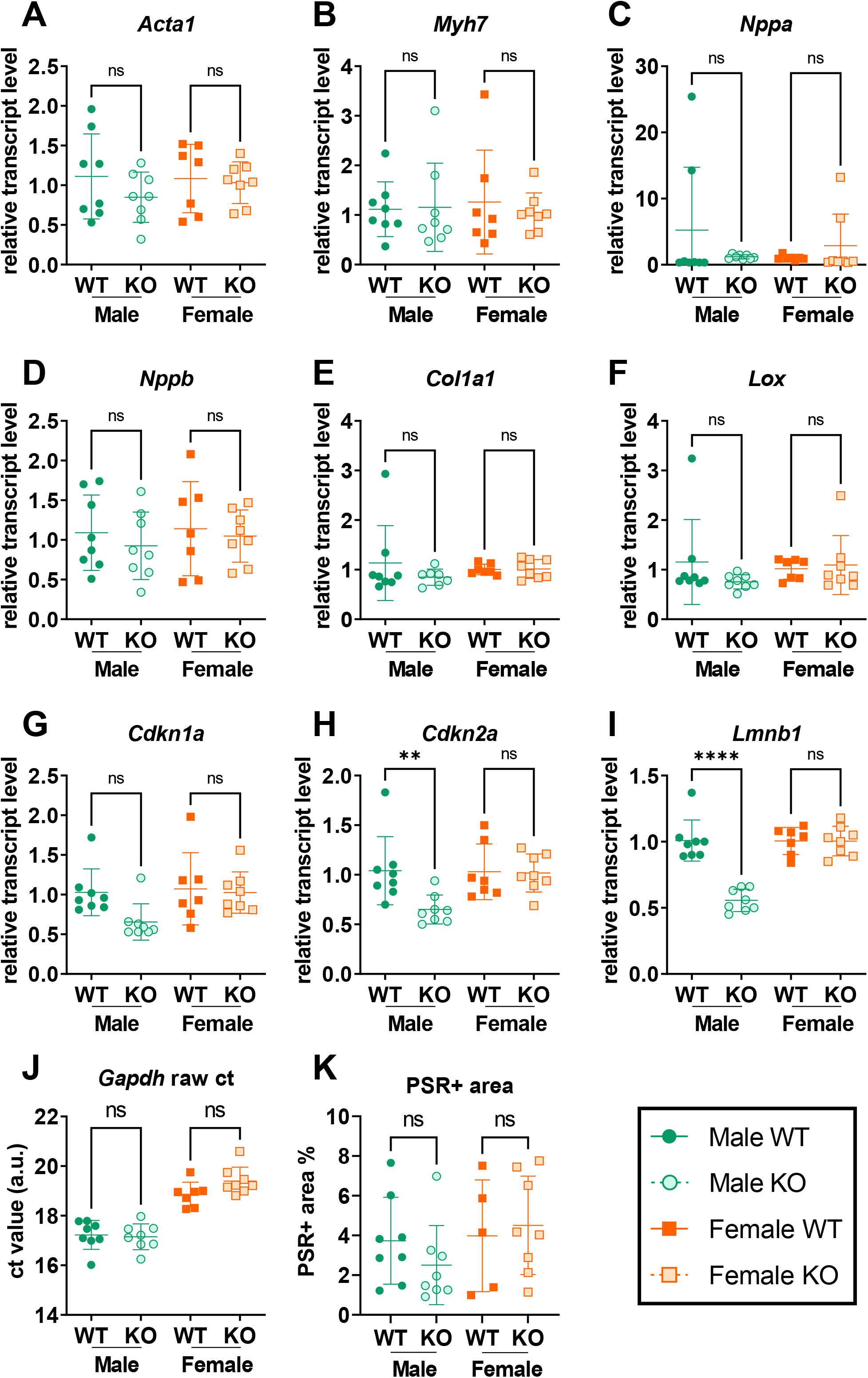
No transcriptional nor histological correlates for echocardiography phenotype of global *Alms1* knockout. (**A-I**) Transcriptional analysis of hearts at 24 weeks old. Data presents ct values normalised to G*apdh* run in duplex. (**J**) An example of raw *Gapdh* ct values from duplexed reactions. (**K**) Quantification of picrosirius red (PSR) staining by pixel thresholding. Samples analysed from 24 week old male and female *Alms1* knockout mice. Each data point represents an individual animal, with bars representing mean ± sd. Comparison between groups performed using a two-way ANOVA with Tukey’s multiple comparisons test. ** denotes p<0.01. N = 8, 8, 7 and 8 for WT males, KO males, WT females and KO females respectively.

Finally, in order to assess the contribution of different cell populations to the cardiac dysfunction seen in female mice at 23 weeks of age, the cardiac phenotype of mice in which *Alms1* KO was driven by the *Pdgfrα* promoter, a marker of mesenchymal stem cells, was evaluated. As would be expected from loss of *Alms1* in cardiac fibroblasts and ∼18% cardiomyocytes, *Alms1* myocardial transcript levels were significantly decreased but not completely lost in hearts of female MSC-specific *Alms1* KO mice (**Figure S4A**). Similar to female global *Alms1* KO mice, heart mass was proportionate to body mass in female MSC-specific *Alms1* KO animals (**Figure 4A**), but in contrast to female global *Alms1* KO mice, female MSC-specific *Alms1* KO mice did not show differences in left atrial area, ejection fraction, fractional area change nor fractional shortening on echocardiography (**Figure 4B-E**). A decrease in isovolumic relaxation time did remain in MSC-specific *Alms1* KO mice (**Figure 4F**), but all other echocardiographic indices were unchanged (**Figure S4C-J**) except myocardial performance index, which was decreased (**Figure S4G**).

**Figure 4.**
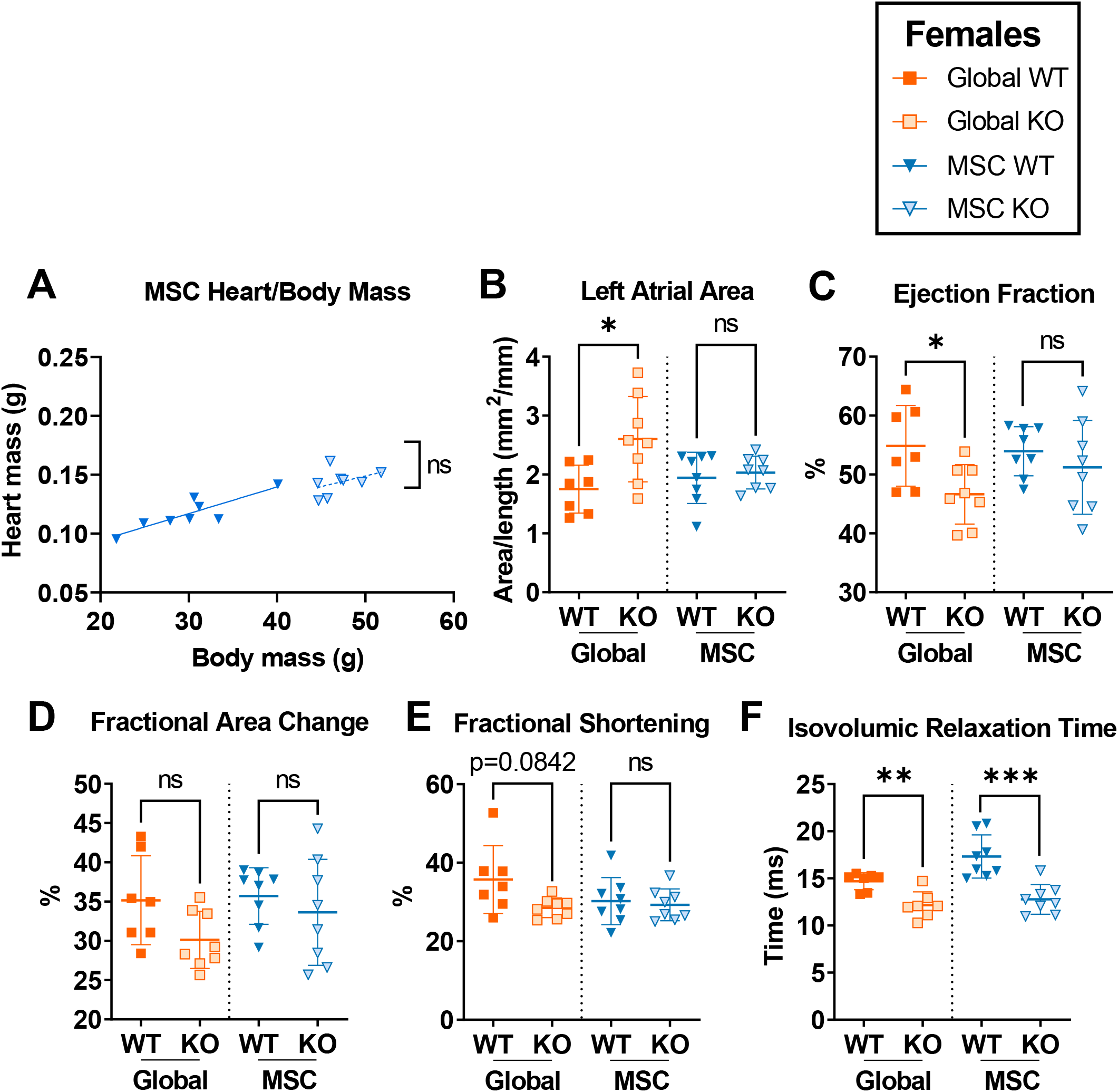
Mesenchymal stem cell-specific *Alms1* knockout in female mice does not recapitulate the phenotype of global *Alms1* knockout. All global knockout (KO) data are repeated from figure 2 for comparison to mesenchymal stem cell- (MSC-) specific *Alms1* KO. **(A)** Linear regression of heart to body mass at 24 weeks of age (**B-F**) Data obtained from analysis of echocardiography performed at 23 weeks of age. Left atrial area (**B**) normalised to nose-anus length. Each data point represents an individual animal with bars in (**B-F**) representing mean ± sd. Lines in linear regression graph (**A**) represent lines of best fit. Global WT/KO and MSC WT/KO experiments were performed with identical design at different times; this is reflected in the dotted line separating the two cohorts. Comparison between WT and KO in (**B-F**) was undertaken using an unpaired two-tailed Student’s t-test followed by a Bonferroni correction for multiple testing. Comparisons between lines of best fit (**A**) was undertaken using simple linear regression, with square brackets showing comparison of y intercepts. No significant change was seen between gradients. * denotes p<0.05, ** denotes p<0.01 and *** denotes p<0.001. N = 7, 8, 8 and 8 for global WT, global KO, MSC WT and MSC KO respectively.

## Discussion

This study generated two *in vivo* models of AS, namely a new global KO and an MSC-specific *Alms1* KO, both derived from the same *Alms1* floxed parental line. Both were extensively characterised with respect to cardiac structure and function at multiple time points. Despite the prominence of cardiomyopathy in AS, and the high importance assigned to improving understanding of this by patients and families, such analysis has not been reported for any prior *Alms1* KO model. We find that, at 23 weeks of age, global *Alms1* KO female mice do show evidence of systolic dysfunction (reduced ejection fraction, fractional area change and fractional shortening), although neither myocardial performance index nor global longitudinal strain were significantly changed. Diastolic dysfunction was suggested by increased left atrial area and decreased isovolumic relaxation time^25^, but longitudinal strain rate, another indicator of diastolic dysfunction, was unchanged. Despite the echocardiographic abnormalities seen in female global *Alms1* KO mice at 23 weeks of age, no corresponding transcriptional or histological changes were detected in myocardial tissue, including no evidence of increased fibrosis or senescence. Nevertheless our findings collectively do indicate mild combined systolic and diastolic dysfunction, reminiscent of the restrictive cardiomyopathy of AS^26^, at least in adult female *Alms1* KO mice. As age is an important cofactor in development of AS cardiomyopathy, it is plausible that more pronounced cardiac dysfunction may become manifest at more advanced ages.

In contrast to findings in older female KO mice, no phenotype was observed in male *Alms1* KO mice at the same age. This is notable, as males have been found to be more severely affected than females for several mouse models of genetic cardiomyopathy^27,28^. No sexual dimorphism in cardiac pathology in human AS has been reported^7^. Diastolic dysfunction has been associated with non-cardiac drivers such as hypertension, diabetes, and obesity^29^, and we have reported that the line of global *Alms1* KO mice studied here, like other global KO models described previously, is obese and insulin resistant^19^. The difference in these metabolic traits is much more pronounced between female wild-type and KO mice than between male counterparts, and male KO mice are severely insulin resistant but not hyperglycaemic^19^. This implicated metabolic differences as one possible cause of the sexual dimorphism we describe in the *Alms1* KO cardiac phenotype. In keeping with this, Heart Failure with preserved Ejection Fraction (HFpEF), which is associated with diabetes, is more prominent in females than males^30^, while diabetes increases the risk of heart failure twice as much in females as in males^31^. Although no correlation has been found cross sectionally between metabolic dysregulation and cardiac dysfunction in human AS, numbers studied have been very small^32^.

To narrow the search for the cellular origin of the cardiac phenotype of older female global *Alms1* KO mice, we also studied female mice with *Alms1* KO driven by *Pdgfra*-cre. This has been shown to delete *Alms1* in a raft of mesenchymal stem cells, including cardiac fibroadipogenic precursors, while largely sparing cardiomyocytes^18^. In this study the indices of systolic and diastolic dysfunction seen in female global *Alms1* KO mice were not recapitulated by female MSC-specific *Alms1* KO mice. Significantly, these conditional KO mice do also have insulin resistance and diabetes^19^. This is counter to the suggestion that the sexually dimorphic cardiomyopathy in female *Alms1* KO mice is purely metabolically mediated, instead indicating that cardiomyocyte-autonomous *Alms1* deficiency plays an important, and perhaps dominant role. Cardiomyocyte-specific *Alms1* KO could be used to confirm this in future.

A limitation of this study is that blood pressure was not measured, but *Alms1* KO rats are reported to be hypertensive, attributed to altered tubular trafficking of the Na-K-Cl channel, and hypertension is observed in around 30% of patients with AS^2,14,32^. It is thus highly plausible that our new KO line, too, features hypertension, and future assessment of this in this model is warranted.

This study did not find evidence of cardiomyopathy in *Alms1* KO mice at P15, at odds both with the previous demonstration in *Alms1^GT/GT^*mice of increased heart/body mass ratio and persistent cardiomyocyte mitosis, and with mitogenic cardiomyopathy in four infants with biallelic *ALMS1* mutations^3^. Given the good concordance of several important phenotypic features of AS among different *Alms1* KO mice models, it was surprising that this early cardiac phenotype could not be replicated. The discrepancy between studies of AS-related infantile cardiomyopathy may relate to genetic background, as the prior study used mice with a mixed 129/C57BL6/J background. Genetic background has been shown to influence other murine cardiac phenotypes^33–35^, while the incomplete penetrance (43%) of infantile cardiomyopathy in AS also argues that environmental or background genetic factors may play a role in human expression of AS cardiomyopathy. Cardiomyocyte binucleation is typically complete by p10 in WT mice^3^, and it remains possible that a difference in trajectory of cardiomyocyte maturation would have been discerned if an earlier time point had been studied. Indeed, infantile cardiomyopathy in patients with AS commonly spontaneously recovers, in keeping with a self-limiting alteration of maturation kinetics. Future studies of *Alms1* KO mice incorporating multiple early postnatal time-points may thus be informative.

In toto, although we detect early cardiac abnormalities in 23 week old female global *Alms1* KO mice, this study demonstrates that mice do not faithfully replicate the severe biphasic cardiomyopathy common in human AS. This may reflect a fundamental inter-species difference in the consequences of *Alms1* loss, undermining the future utility of mouse models for its study. Alternatively, it may be revealing about the relative contributions of different pathogenic mechanisms to heart failure in human AS. Two pertinent differences between C57BL6/N mice and humans are that that the mice are neither extremely fibrosis-^36^ nor atherosclerosis-prone. The absence of a severe cardiac phenotype in mice could thus be interpreted as strengthening the case that accelerated atherosclerosis, and/or pathological abnormalities in remodelling in the face of ischaemia, and/or excess fibrosis are the dominant drivers of adult heart failure in AS.

Several other murine models of cardiomyopathy model human disease poorly (e.g.^37–39^). Several measures are worthy of consideration in future to increase the value of the *Alms1* KO mouse as a cardiac disease model. These include increasing the age of the mice studied, as age is an important factor in development of diastolic dysfunction^29,34^, or using pharmacological stressors such as angiotensin II or adrenergic agonists^40^ to ‘unmask’ cardiac dysfunction. More specific options to gain insights into AS heart failure pathogenesis include examining *Alms1* KO on either an atherosclerosis-prone (e.g. *Apoe^-/-^* or *Pcsk9^-/-^*) or fibrosis-prone (S129S6) genetic background. Our findings, however, underline the value of alternative experimental models of the cardiac complications of AS, including for example the induced pluripotent stem cell-derived cardiomyocytes. Continuing to optimise models of cardiac complications of AS will be crucial not only for unpicking and more effectively targeting devastating complications of an ultra-rare disease, but may also yield prismatic insights into the role of complex multi-morbidities (e.g. diabetes, obesity, kidney and liver impairment) in more common conditions such as HFpEF.

## Author Contributions

**EM:** conceptualisation, methodology, formal analysis, investigation, writing - original draft, writing - review & editing, visualization; **IL**: supervision, writing - review & editing; **AT:** methodology, formal analysis, investigation; **XW**: investigation; **KG:** methodology, writing - original draft, writing - review & editing; **GG:** writing - original draft, writing - review & editing, supervision; **RKS:** conceptualisation, methodology, resources, writing - original draft, writing - review & editing, supervision, funding acquisition.

