## Supplementary Information for "Female Alms1-deficient mice develop echocardiographic features of adult but not infantile Alström Syndrome cardiomyopathy"

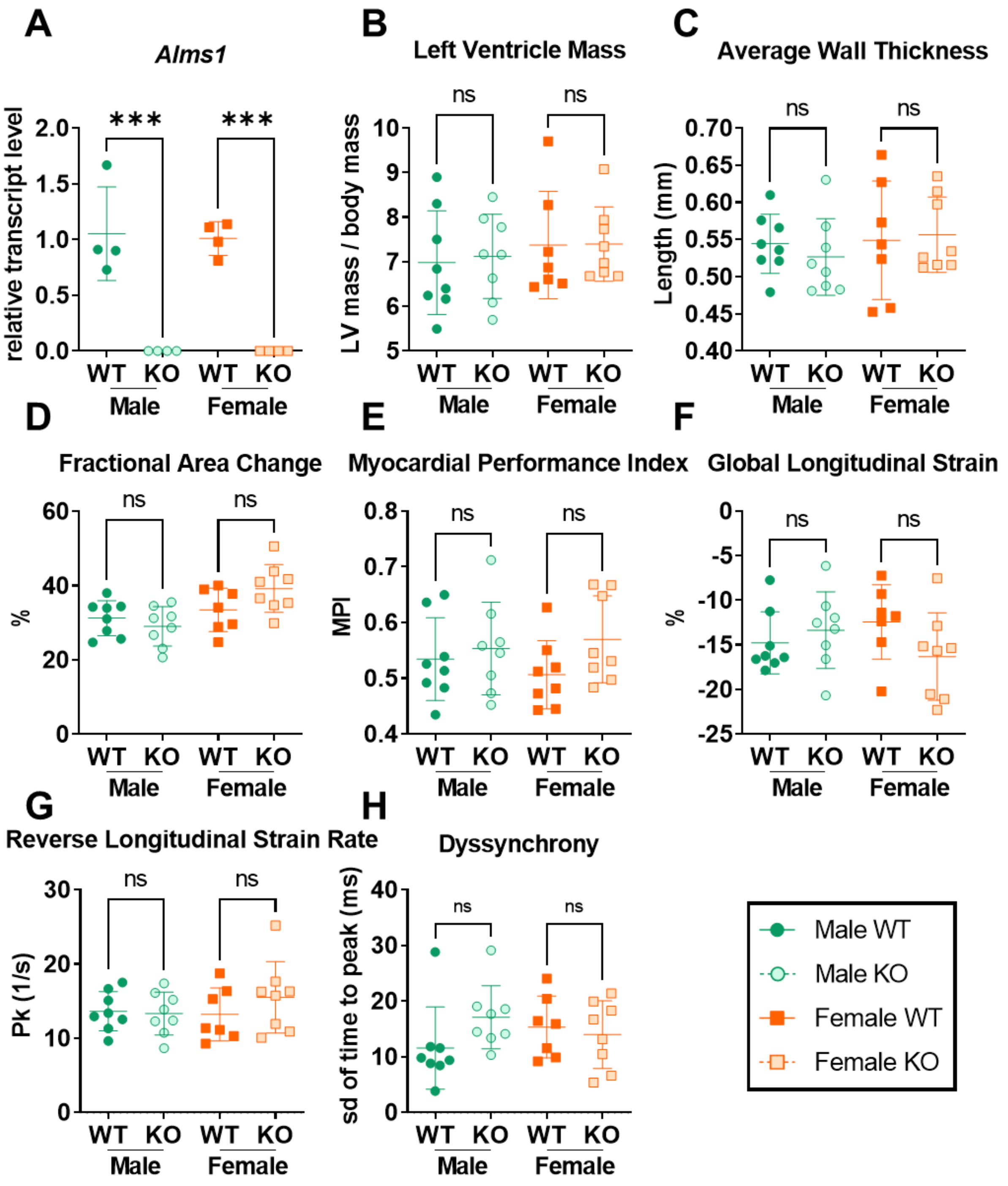


**Figure S1. Neither male nor female global *Alms1* knockout mice exhibit an echocardiographic phenotype at post-natal day 15.** (**A**) qPCR confirmation of *Alms1* loss in heart tissue of global *Alms1* knockout (KO) mice with a Taqman probe for the 6-7 exon junction of *Alms1* **(B-H)** Echocardiography data. Left ventricle mass values (**B**) are normalised to total body mass. (**A**) Data presents ct values normalised to *Gapdh* run in duplex. Each data point represents an individual animal with bars representing mean ± sd. Comparison between groups in performed using two-way ANOVA with Tukey's multiple comparisons test. For (**A**) N = 4/group. For (**B-H**) N = 8, 8, 7 and 8 for WT males, KO males, WT females and KO females respectively.

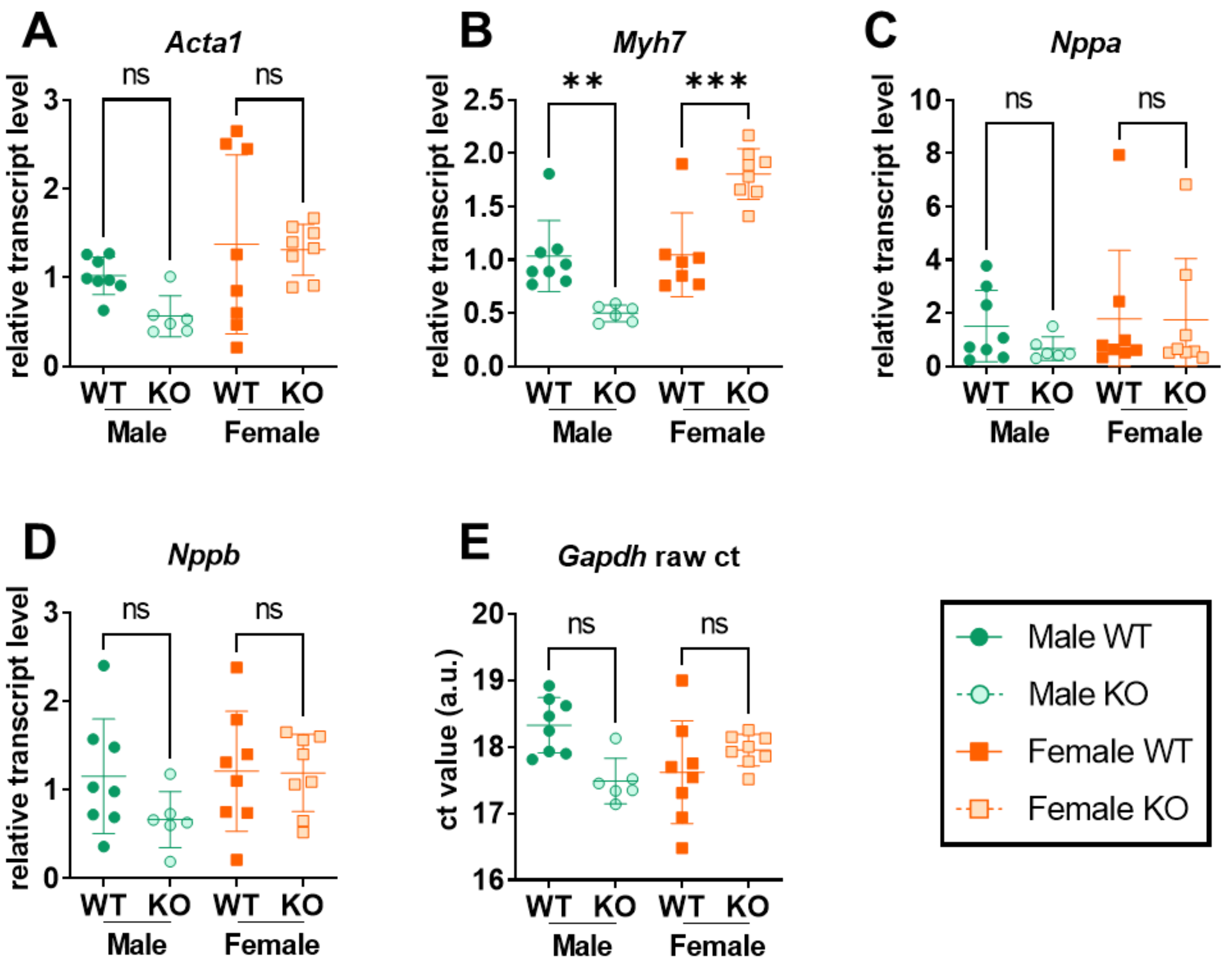


**Figure S2. Neither male nor female global *Alms1* knockout mice exhibit a cardiac transcriptional phenotype at post-natal day 15.** (**A-D**) qPCR evaluation of heart tissue for typical markers of cardiomyopathy. Data presents ct values normalised to *Gapdh* run in duplex. (**E**) An example of raw *Gapdh* ct values from duplexed reactions. Each data point represents an individual animal with bars representing mean ± sd. Comparison between groups in performed using two-way ANOVA with Tukey's multiple comparisons test. N = 8, 8, 7 and 8 for WT males, KO males, WT females and KO females respectively.

Female Alms1-deficient mice develop echocardiographic features of adult but not infantile Alström Syndrome cardiomyopathy

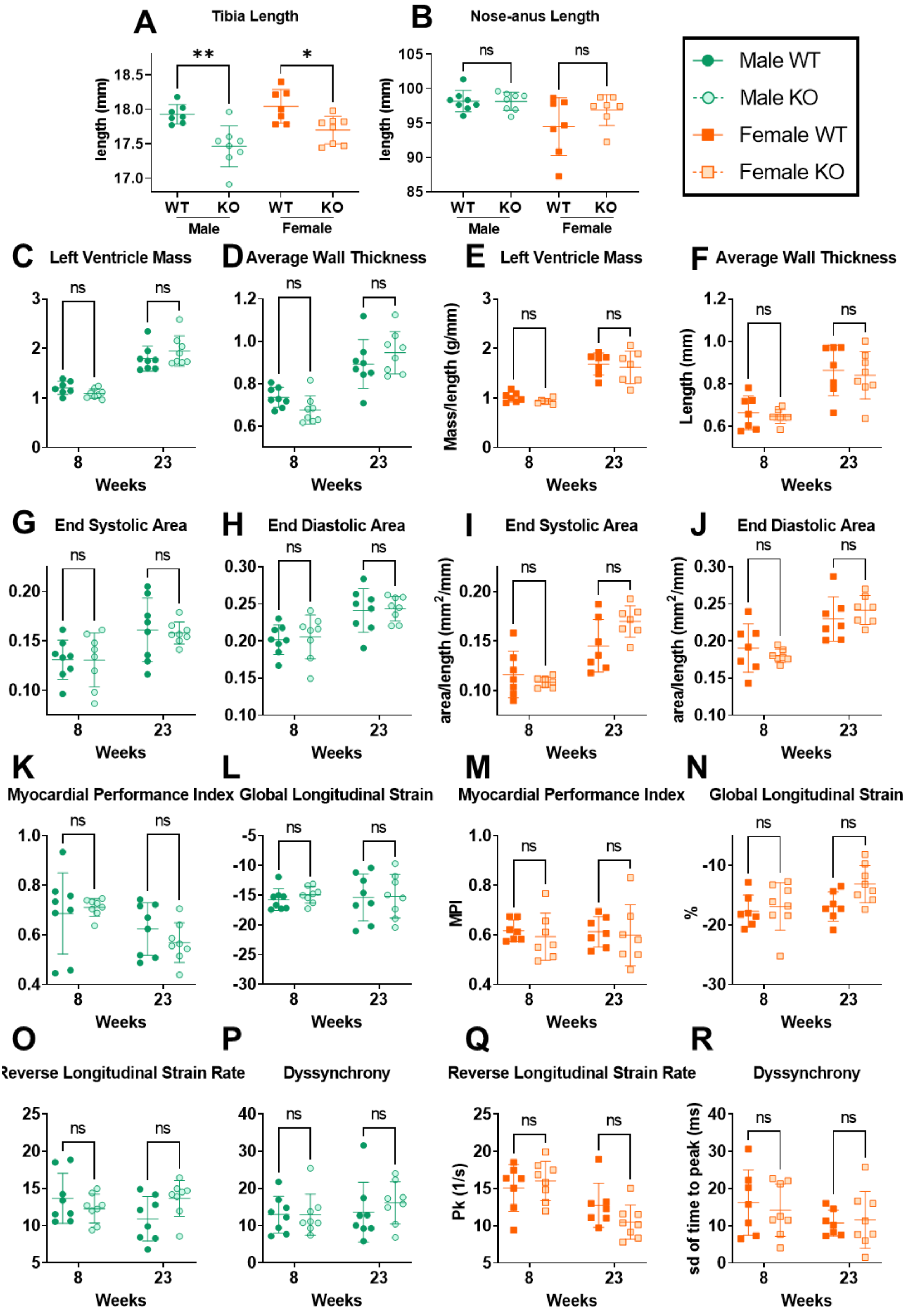

### Female Alms1-deficient mice develop echocardiographic features of adult but not infantile Alström Syndrome cardiomyopathy

**Figure S3. Systolic and diastolic dysfunction develops in female but not male global *Alms1* knockout mice with age. (A,B)** Evaluation of parameters for normalisation of echocardiography area and mass values. (**A**) Tibia length measured following cull at 24 weeks. (**B**) Nose-anus length measured in anaesthetised animals immediately following echo at 23 weeks. (**C-R**) Echocardiography parameters measured at 8 and 23 weeks of age. Each data point represents an individual animal with bars representing mean ± sd. Left ventricle mass and area values (**C,E,G-J**) are normalised to nose-anus length. Comparison between groups in (**A,B**) performed using two-way ANOVA with Tukey's multiple comparisons test. Comparison between groups (**C-R**) performed using a two-way ANOVA with Šídák's multiple comparisons test. * denotes $p<0.05$ and ** denotes $p<0.01$. N = 8, 8, 7 and 8 for WT males, KO males, WT females and KO females respectively

Female Alms1-deficient mice develop echocardiographic features of adult but not infantile Alström Syndrome cardiomyopathy

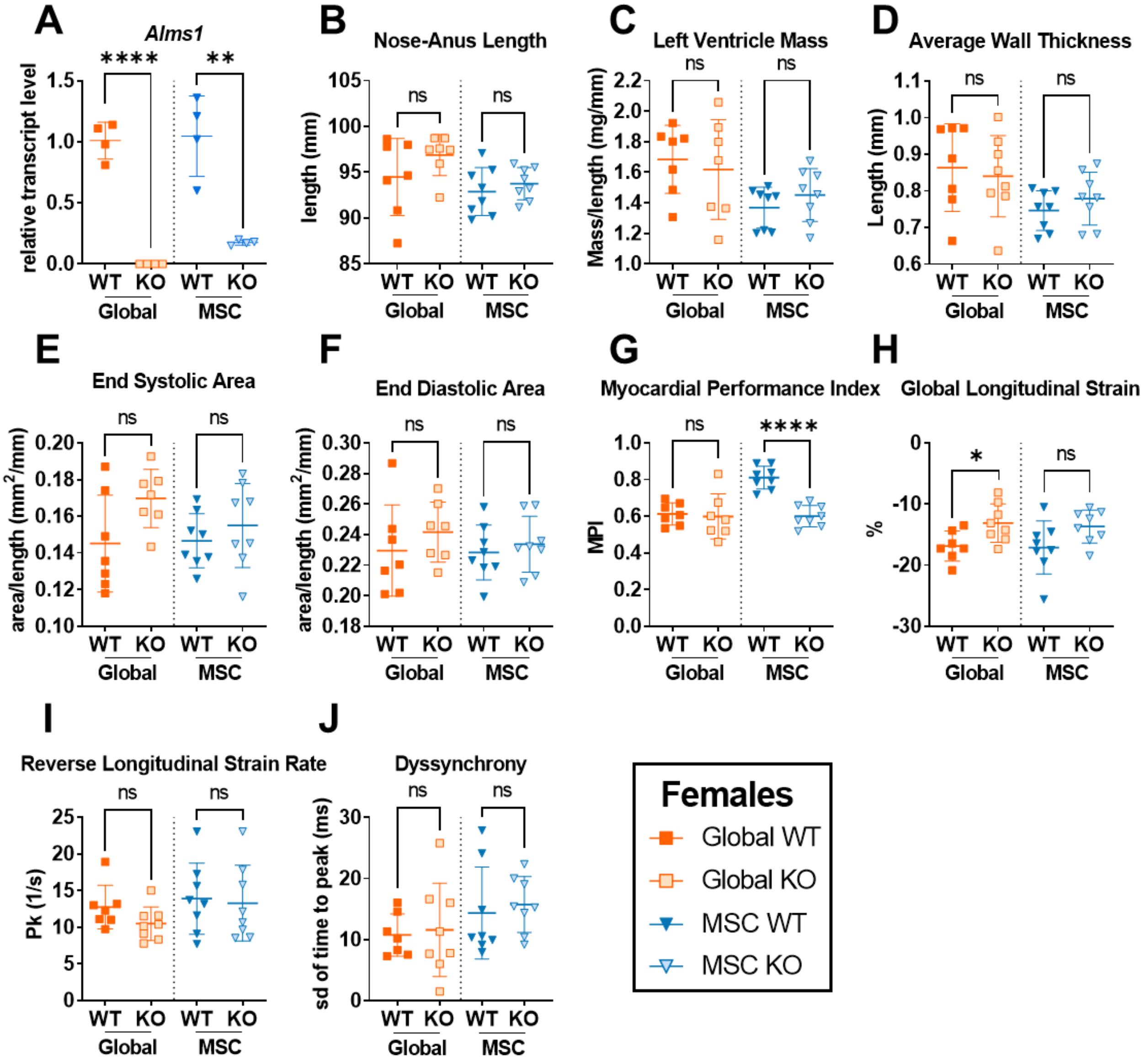


**Figure S4. Mesenchymal stem cell-specific *Alms1* knockout does not recapitulate the phenotype of global *Alms1* knockout.** All global KO data repeated from Figure 2 and Figure S2 for comparison to MSC-specific *Alms1* KO. (**A**) qPCR confirmation of partial *Alms1* loss in heart tissue of MSC-specific Alms1 KO mice with a Taqman probe for the 6-7 exon junction of *Alms1*. (**B**) Nose to anus length of females at 23 weeks of age. (**C-J**) Data obtained from analysis of echocardiography performed on female animals at 23 weeks of age. Mass and area values (**C,E,F**) normalised to nose-anus length. Each data point represents an individual animal with bars representing mean ± sd. Global WT/KO and MSC WT/KO experiments were performed with identical design at different times; this is reflected in the dotted line separating the two cohorts. Comparison between WT and KO was performed using an unpaired two-tailed Student's t-test followed by a Bonferroni correction for multiple testing. * denotes p<0.05, ** denotes p<0.01, *** denotes p<0.001 and **** denotes p<0.0001. (**A**) N = 4/group (**B-J**) N = 7, 8, 8 and 8 for female global WT, global KO, MSC WT and MSC KO respectively.

#### Supplementary Methods

*Electrocardiographic analysis*

Analysis of echocardiographic data was performed using Visualsonics Vevo LAB 5.71 software (FUJIFILM VisualSonics). End systolic and end diastolic area, ejection fraction, fractional area change, left ventricle mass and average wall thickness were calculated from the PSLA EKV. End systolic and end diastolic area and ejection fraction were calculated using the automated artificial-intelligence software AutoLV (Grune *et al.*, 2019). Fractional area change, left ventricle mass and average wall thickness were calculated after manual tracing of the contours of PSLA epicardial and endocardial area and axis length at systole and diastole, as indicated by AutoLV. Fractional shortening was calculated from M-mode images using AutoLV. Modified right side PSLA EKV was used to manually measure left atrium area, choosing the time frame immediately after the mitral valve closure. Isovolumic relaxation time (IVRT), isovolumic relaxation time (IVCT) and ejection time (ET) were all calculated by manual analysis of Doppler; three consecutive annotations for each index were made. IVRT measures the time between aortic valve closing and the mitral valve opening. Myocardial performance index (MPI) was calculated as the sum of IVRT and IVCT divided by ET. Finally, the 'Vevo Strain' function of the Vevo LAB software was used to analyse PSLA EKV, generating values for global longitudinal strain (GLS), reverse longitudinal strain rate (rLSR) and ventricular dyssynchrony. Ventricular dyssychrony was calculated as the standard deviation of strain across the 6 panels generated by 'Vevo Strain'.

Female Alms1-deficient mice develop echocardiographic features of adult but not infantile Alström Syndrome cardiomyopathy

*Tissue studies*

Mice were culled by cervical dislocation under isoflurane anaesthesia. Dissected hearts were rinsed in phosphate-buffered saline (PBS), and weighed after blotting of excess fluid. Whole neonatal hearts were fixed in 4% paraformaldehyde or snap-frozen in liquid nitrogen. The bottom 1/5$^{th}$ (apex) of adult hearts was fixed in 4% PFA, and the middle 2/5ths (ventricles) snap-frozen in liquid nitrogen. Adult tibia were also collected.

For histological analysis, fixed cardiac tissue was paraffin-embedded and 5μm sections cut. For picrosirius red (PSR) staining slides were dewaxed in xylene, rehydrated in decreasing concentrations of ethanol and washed in water before incubation in direct red picric acid solution for 2 hours, washing, dehydrating in ethanol, clearing in xylene, and mounting. Immunofluorescence staining of neonatal heart sections was performed using the Leica BOND III immunostainer robot at room temperature, with reagents detailed in **Supplementary Table 1**. Sequential staining for histone H3, cardiac troponin (cTnT) and wheat germ agglutinin (WGA) was performed, with antibodies and conditions used detailed in **Supplementary Table 2**. Washing with TBS-T buffer (composition) was performed between steps. For histone H3, antigen retrieval was by incubation in Bond Epitope Retrieval ER1 Solution (Leica Microsystems) for 20 minutes before peroxide blocking, blocking in diluted goat serum diluted 1:5 for 10 minutes. Anti-histone H3 antibody was incubated for 60 minutes before incubation with Goat Anti-Rabbit IgG HRP-conjugated secondary antibody for 30 minutes. Finally tyramide signal amplification was performed with addition of tyramide substrate FITC green opal 520. Next cTnT staining was performed. Firstly antigen retrieval was performed by incubation in Bond Epitope Retrieval ER1 Solution for 10 minutes. Peroxide blocking was performed for 10 minutes followed by serum blocking for 10 minutes using blocking solution from the Mouse on Mouse Polymer IHC Kit (Abcam). Incubation of anti-cTnT was performed for 60 minutes before addition of secondary polymer from the Mouse on Mouse Polymer IHC Kit (Abcam) for 30 minutes. Finally, the tyramide substrate blue CY5 opal 650 was added. Next WGA staining was performed. Antigen retrieval was performed by incubation in Bond Epitope Retrieval ER1 Solution for 10 minutes. Peroxide blocking was then performed for 10 minutes before serum blocking in goat serum diluted 1:5 for 10 minutes. Rhodamine-conjugated WGA diluted 1:75 was then incubated for 60 minutes. Finally nuclear counter-staining was performed by incubation with DAPI, diluted 1:1000. All histological slides were imaged using a Zeiss Axioscan.Z1 with Zen2.6 software.

*Gene expression analysis*

RNA was extracted from snap-frozen heart tissue using the Qiagen RNeasy Fibrous Tissue Mini Kit after homogenisation in 2mL tubes containing 2.8mm ceramic beads stored on dry ice, using the Omni Bead Ruptor 24 Elite with the pre chilled Omni Cryo unit filled with dry ice. 10µL of RLT buffer was added per 1mg of heart tissue. 300µL homogenate was used for RNA extraction. After elution, RNA concentration was measured using the NanoDrop ONE before dilution to 100ng/µL in nuclease-free water. Reverse transcription was performed using the High Capacity cDNA Reverse Transcription Kit (Applied Biosystems) in the Eppendorf Mastercycler X50s, using 1000ng per reaction. cDNA solution was then diluted 1 in 4 with nuclease-free water. Control reactions without reverse transcriptase were performed alongside all experimental reactions.

Real-time quantitative PCR (RT-qPCR) was performed using TaqMan reagents on a LightCycler® 480 Instrument II (Roche) in duplex with minor groove binder (MGB) probes (Hein and Bodendorf, 2007). *Gapdh* was evaluated as a housekeeping control gene for normalisation using a 2'-chloro-7'phenyl-1,4-dichloro-6-carboxy-fluorescein (VIC)-coupled probe. Primer efficiency was calculated by dilution standard curve prior to experimental reactions. All reactions were run in triplicate, and RT- and non-template controls were run on the same plate. Crossing point (Cp) values were calculated using LightCycler 480 software using the Abs quant / 2nd derivative max function. Cp values for the gene of interest were normalised to duplexed *Gapdh* Cp values after adjusting for primer efficiency, as first described by Pfaffl (Pfaffl, 2001). Raw Cp values for the gene of interest and *Gapdh* were also visualised in all cases (e.g. **Figure 3J, S2E**). Taqman primer and probe mixes used for qPCR were purchased from Life Technologies and are listed in **Supplementary Table 3**.

*Supplementary Tables*

| **Item** | **Catalog No.** | **Company** |
|---|---|---|
| Bond Epitope Retrieval ER1 Solution | AR9961 | Leica Biosystems |
| Bond Epitope Retrieval ER2 Solution | AR9640 | Leica Biosystems |
| Bond Wash Solution | AR9590 | Leica Biosystems |
| Bond Polymer Refine Detection Kit | DS9800 | Leica Biosystems |
| Normal Goat Serum | ab7481 | Abcam |
| Mouse on Mouse Polymer IHC Kit | ab269452 | Abcam |
| Rhodamine-conjugated Wheat Germ Agglutinin | RL-1022 | Vector Laboratories |
| DAPI | D3571 | Life Technologies |
| 520 Green Opal reagent pack | FP1487001KT | Akoya |
| 650 Blue opal reagent pack | FP1496001KT | Akoya |

**Supplementary Table 1.** Reagents used for immunohistochemistry.

| **Antibody** | **Catalog No.** | **Species raised in** | **Company** | **Dilution** |
|---|---|---|---|---|
| anti-cTnT (cardiac troponin) | MA512960 | Mouse | Invitrogen, USA | 1:500 |
| Anti-Histone H3 (phospho S10) | ab5176 | Rabbit | Abcam, UK | 1:400 |
| Goat F(ab) Anti-Rabbit IgG H&L (HRP) | ab7171 | Goat | Abcam, UK | 1:500 |

**Supplementary Table 2.** Antibodies used for immunohistochemistry.

| **Reagent** | **Gene target** | **Taqman Probe ID** | **Catalog No.** |
|---|---|---|---|
| Mouse GAPD (GAPDH) Endogenous Control (VIC™/MGB probe, primer limited) | *Gapdh* | Mm99999915_g1 | 4352339E |
| TaqMan™ Gene Expression Assay (FAM) | *Alms1* exon 6-7 | Mm01189441_m1 | 4351372 |
| TaqMan™ Gene Expression Assay (FAM) | *Acta1* | Mm00808218_g1 | 4331182 |
| TaqMan™ Gene Expression Assay (FAM) | *Myh7* | Mm00600555_m1 | 4331182 |
| TaqMan™ Gene Expression Assay (FAM) | *Nppa* | Mm01255748_g1 | 4331182 |
| TaqMan™ Gene Expression Assay (FAM) | *Nppb* | Mm01255770_g1 | 4331182 |
| TaqMan™ Gene Expression Assay (FAM) | *Col1a1* | Mm00801666_g1 | 4331182 |
| TaqMan™ Gene Expression Assay (FAM) | *Lox* | Mm00495386_m1 | 4331182 |
| TaqMan™ Gene Expression Assay (FAM) | *Cdkn1a* | Mm04205640_g1 | 4331182 |
| TaqMan™ Gene Expression Assay (FAM) | *Cdkn2a* | Mm00494449_m1 | 4331182 |
| TaqMan™ Gene Expression Assay (FAM) | *Lmnb1* | Mm00521949_m1 | 4331182 |

**Supplementary Table 3.** TaqMan primer/probe mixes.
